# The shared genetic architecture and evolution of human language and musical rhythm

**DOI:** 10.1101/2023.11.01.564908

**Authors:** Gökberk Alagöz, Else Eising, Yasmina Mekki, Giacomo Bignardi, Pierre Fontanillas, 23andMe Research Team, Michel G. Nivard, Michelle Luciano, Nancy J. Cox, Simon E. Fisher, Reyna L. Gordon

**Affiliations:** Language and Genetics Department, Max Planck Institute for Psycholinguistics, 6500 AH Nijmegen, The Netherlands; Department of Otolaryngology - Head & Neck Surgery, Vanderbilt University Medical Center, Nashville, TN, USA; Vanderbilt Genetics Institute, Vanderbilt University Medical Center, Nashville, TN, USA; Max Planck School of Cognition, Leipzig, Germany; 23andMe, Inc., Sunnyvale, CA, USA; Department of Biological Psychology, Vrije Universiteit, Amsterdam, the Netherlands; Department of Psychology, University of Edinburgh, Edinburgh, UK; Donders Institute for Brain, Cognition and Behaviour, Radboud University, 6500 HB Nijmegen, The Netherlands; Vanderbilt Brain Institute, Vanderbilt University, Nashville, TN, USA; The Curb Center, Vanderbilt University, Nashville, TN, USA

## Abstract

Rhythm and language-related traits are phenotypically correlated, but their genetic overlap is largely unknown. Here, we leveraged two large-scale genome-wide association studies performed to shed light on the shared genetics of rhythm (N=606,825) and dyslexia (N=1,138,870). Our results reveal an intricate shared genetic and neurobiological architecture, and lay groundwork for resolving longstanding debates about the potential co-evolution of human language and musical traits.

## Main text

The human brain has evolved intricate neural circuitry to process complex communicative signals and behaviours, including speech and music, and the extent of biological overlap between these facets is an important question for the field of neurobiology. Individual differences in rhythm-related skills (e.g., beat synchronisation, rhythm perception and production, metrical perception) are correlated with variability in a range of language-related skills (e.g., word recognition, spelling, phonological awareness), implicating potentially shared underlying neural and genetic architectures^1^. In particular, individuals with rhythm impairment have been suggested to show higher predisposition to language-related difficulties such as dyslexia and developmental language disorder (Atypical Rhythm Risk Hypothesis, ARRH)^2^. Given that disorders of language and reading can have long-term health impacts, identifying genetic factors that they share with rhythm impairment may enhance future screening capabilities. Moreover, basic science concerning the biological substrates of these fundamental human traits will be informed by new approaches to their potentially shared genetic architecture.

The evolution of rhythm-related traits has been hypothesized to be linked to multiple facets of human communication, including parent-child bonding, social or group cohesion, and aspects of speech/language^3,4^. To address prominent theories on the co-evolution of phonological skill development and rhythm in humans^5^, evidence to date has been taken largely from psychology, neuroscience, and cross-species comparisons rather than genetics^6,7^. We hypothesise that identifying the shared genetic architecture between rhythm- and language-related disorders, and probing the evolutionary past of the implicated genomic regions, can help reveal neural and biological characteristics of our species which made rhythm and language an asset to human development and behaviour.

Our work built on two recent genome-wide association studies (GWAS) that represent by far the most well-powered genetic investigations of rhythm-/language-relevant traits to date, one for musical rhythm (beat synchronisation, hereafter referred to as *rhythm*; “Can you clap in time with a musical beat?”, N_cases_(Yes)=555,660, N_controls_(No)=51,165)^8^ and the other for dyslexia (developmental reading/spelling difficulties; “Have you been diagnosed with dyslexia?”, N_cases_(Yes)=51,800, N_controls_(No)=1,087,070)^9^, both performed on 23andMe, Inc. Research Cohort in individuals of European ancestry, and both classified as binary traits. We used the dyslexia GWAS as a proxy for the genetic underpinnings of language and reading-related aspects of human communication, as dyslexia often co-occurs with a number of speech/language disorders^10,11,12,13^. Beat synchronisation GWAS was used as a proxy for musical rhythm skills, as beat perception and synchronisation are considered to be important features of musical experiences in present-day humans^14,15^. We applied a three-stage analytic pipeline to investigate shared genetics and biology: i) Genome-wide genetic correlations between rhythm and dyslexia (as well as other language-related traits) using linkage disequilibrium score regression (LDSC) ^16^, ii) multivariate GWAS (mvGWAS) of rhythm impairment and dyslexia using Genomic Structural Equation Modelling (SEM)^17^, iii) post-mvGWAS analyses of the shared genomic infrastructure as windows into its evolution and biology (Fig. 1A).

**Fig. 1:**
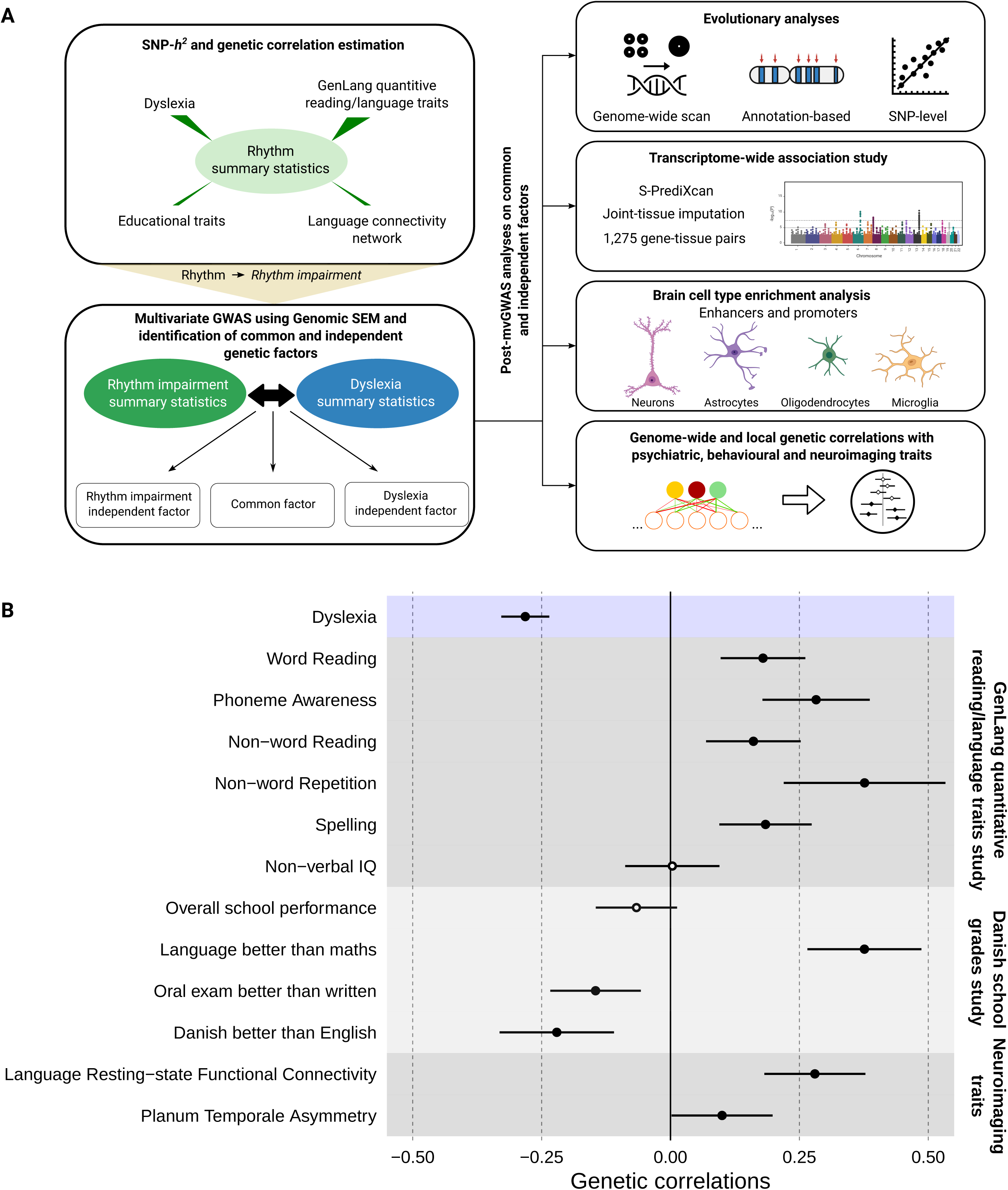
Study design and genetic correlations between rhythm and language-/reading- related traits. **(A)** Flow chart shows analyses performed in our study. SNP-*h*^2^ and genetic correlations were estimated using LDSC. Effect directions in the rhythm GWAS summary statistics were flipped to obtain a proxy to probe rhythm impairment. Genomic SEM was used to identify common and independent genetic factors of rhythm impairment and dyslexia. As for post mvGWAS analyses, we adopted various methods including LDSC partitioned heritability, GCTB SBayesS, LAVA, and manual SNP-lookups. **(B)** Genetic correlations between rhythm and a set of language- and reading-related traits. Significant genetic correlations were indicated by full circles. Error bars correspond to standard errors.

In the first stage, we estimated genetic correlations between rhythm and dyslexia, as well as quantitative measures of language/reading performance^18^, educational traits^19^, and brain-language related endophenotypes^20,21^ by using LDSC^16^. We found moderate but significant genetic correlations between rhythm and dyslexia (*r_g_*(SE)=-0.28(0.02), P_FDR_=2.05×10^−31^), five quantitative language/reading measures, three educational traits, and two language-relevant neuroimaging endophenotypes (Fig. 1B, Table S1). In contrast there were negligible and non- significant genetic correlations with non-verbal IQ (*r_g_*(SE)=-0.004(0.047), P_FDR_=0.94) and overall school performance (*r_g_*(SE)=-0.066(0.040), P_FDR_=0.11) (Fig. 1B, Table S1). Thus, rhythm is genetically correlated not only with dyslexia, but also multiple language-related phenotypes including word and non-word reading, non-word repetition, phoneme awareness, having better language skills than mathematics, and language resting-state functional connectivity (|*r_g_|* median=0.184, range=0.004-0.376), providing empirical genetic evidence for the ARRH. The absence of significant genetic correlations between rhythm and cognitive traits such as non- verbal IQ and overall school performance provide evidence that genetic sharing between rhythm and dyslexia is not driven by general cognition. These results represent the first direct empirical support for a shared genetic architecture underlying previously observed phenotypic correlations between rhythm and language-related traits^1^ such as dyslexia (Pearson correlation=-0.04[-0.05;- 0.04], t=-25.96, df=363285, P<2.2×10^−16^).

Given that dyslexia is a neurodevelopmental disorder with effects particularly apparent in the written language domain (evident from reading and/or spelling difficulties)^9^, and that other work has shown rhythm impairments associated with dyslexia^10,11,12,13^, we expect it to be genetically and phenotypically linked to impairment in rhythm (hereafter referred to as *rhythm impairment*) rather than rhythm ability. (This expectation is supported by the negative sign of the genetic correlation observed in the first stage of our pipeline above.) Thus, we reversed the effect directions in the binary rhythm GWAS summary statistics in order to align genetic effect directions for rhythm- and reading-impairments. We then performed a mvGWAS on the rhythm impairment and dyslexia GWASs to probe the validity of ARRH at the genetic level, using a bivariate extension of Genomic SEM^17^ that we developed (see Methods). This allowed us to tease apart the genetic effects shared between rhythm impairment and dyslexia from those that are unique to each. We specified a measurement model with a shared genetic factor (F_gRI-D_), which recaptured the genetic correlation between two traits (σ^2^_FgRI-D_(SE)=0.28(0.03)). Similar to Grotzinger et al.^22^, we then applied both the Common Pathway Model (CPM), which regresses single-nucleotide polymorphisms (SNPs) from F_gRI-D_ (Fig. S1), and the Independent Pathways Model solution (IPM), which regresses SNPs directly onto the genetic components of the two traits (Fig. S1). We were thus able to obtain a quantitative per-SNP score quantifying the extent to which any given SNP influences rhythm impairment or dyslexia independent from F _gRI-D_, that is the bivariate genetic heterogeneity (Q_b_).

Our mvGWAS analysis with the CPM resulted in a new set of summary statistics representing the genetic overlap between rhythm impairment and dyslexia, and identified 18 genome-wide significant (P<5×10^−8^) loci associated with F_gRI-D_ (Fig. 2A, Table S2) after genomic control (GC) correction (Fig. S2). We estimated the SNP-heritability of F_gRI-D_ as 13% (SE=0.005) by using LDSC^16^. The strongest mvGWAS signal came from the SNP rs28576629 (P=3.79×10^−14^) on chromosome 3 (Fig. 2A), an intronic variant in *PPP2R3A*, a gene encoding a regulatory subunit of protein phosphatase 2^23^. We validated the Genomic SEM CPM results using two additional mvGWAS methods: 1) N-weighted Genome-Wide Association Meta-Analysis (GWAMA)^24^, and 2) Cross-Phenotype Association Analysis (CPASSOC)^25^. Both methods captured highly similar genomic architectures to the one captured by the CPM (Fig. S3), confirming that the shared genetics of rhythm impairment and dyslexia could be identified consistently regardless of analytic tool. The IPM resulted in two new sets of summary statistics capturing the genetic factors of rhythm impairment and dyslexia that are independent from F_gRI-D_, so-called independent factors (Fig. S4). We used the IPM results to obtain Q_b_ and mapped the per-SNP Q_b_ scores onto CPM mvGWAS results to dissociate the homogeneous (hereafter referred to as pleiotropic) signals from the signals driven by a single GWAS (Fig 2A). We identified 27 genome-wide significant (P<5×10^−8^) heterogeneous loci in the Q_b_ results (Fig. 2A, Table S3), and two of these loci are co-localized with two CPM signals on chromosome 20 (30,690,943- 31,189,993 and 47,514,881-47,821,129), which are mvGWAS signals that are driven by the dyslexia GWAS (Fig. 2A). Our analysis revealed two distinct patterns for CPM mvGWAS hit loci: 16 highly homogeneous (putatively pleiotropic) and two heterogeneous loci indicating different levels of GWAS significance, effect sizes and/or opposite effect directions for these two loci in the rhythm impairment and dyslexia GWASs (see Fig. 2B for representative loci of each type).

**Fig. 2:**
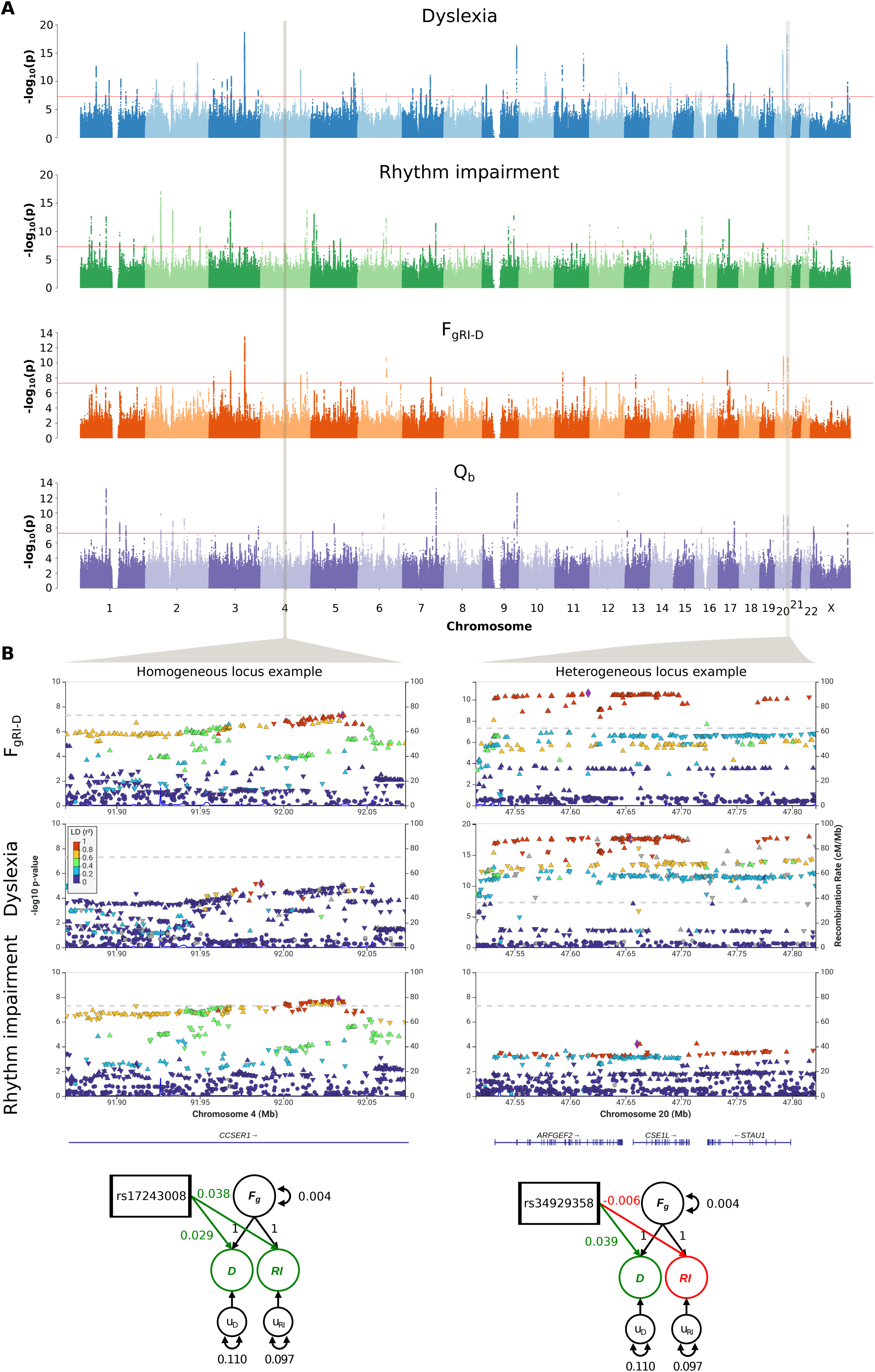
Manhattan plots for univariate and multivariate GWASs and heterogeneity. Examples of highly homogeneous and heterogeneous loci in F_gRI-D_ results. **(A)** Manhattan plots show -log_10_(P) values of dyslexia, rhythm GWASs, F_gRI-D_ mvGWAS and heterogeneity across dyslexia and rhythm impairment. GWAS and mvGWAS results were GC corrected. The red lines correspond to genome-wide significance threshold (P<5×10^−8^). **(B)** LocusZoom plots of example homogeneous and heterogeneous loci, identified according to Q_b_ p-values. SEM diagrams show effect sizes and directions of the selected SNPs for dyslexia and rhythm impairment, reflecting homogeneous vs. heterogeneous architecture of the example loci.

Next, we performed a transcriptome-wide association study (TWAS) using F_gRI-D_ summary statistics, and whole-blood and 13 GTEx brain tissue phenotype weights^26,27^ with S-PrediXcan^28^ (Table S4). Our TWAS analysis identified 1,275 significant (P_FDR_<0.05) gene-tissue pairs, and 315 significant (P_FDR_<0.05) unique genes associated with F_gRI-D_ after FDR correction (Fig. 3A, Table S5). Some of the top significant gene-tissue pairs associated with F_gRI-D_ are *AC072039.2* expression in brain nucleus (Z-score=-7.74, P_FDR_=1.17×10^−9^), *PPP2R3A* expression in cerebellum (Z-score=7.49, P_FDR_=2.43×10^−9^) and putamen (Z-score=7.47, P_FDR_=2.43×10^−9^), and *FOXO3* expression in anterior cingulate cortex (Z-score=6.07, P_FDR_=1.15×10^−5^) (Fig. 3A). Functional enrichment analysis of the significant (P_FDR_<0.05) TWAS genes using PANTHER^29^ did not identify any significant enrichments in Gene Ontology^30,31,32^ and PANTHER GO-Slim^29,30,31,32^ terms after accounting for multiple testing (Tables S6-11). Overall, our S-PrediXcan analysis highlighted 315 unique genes linked to F_gRI-D_, including significant gene-tissue pairs (such as *FOXO3* expression in the anterior cingulate cortex, and *PPP2R3A* expression in the putamen) involving brain regions with known relevance for music processing^33,34^.

**Fig. 3:**
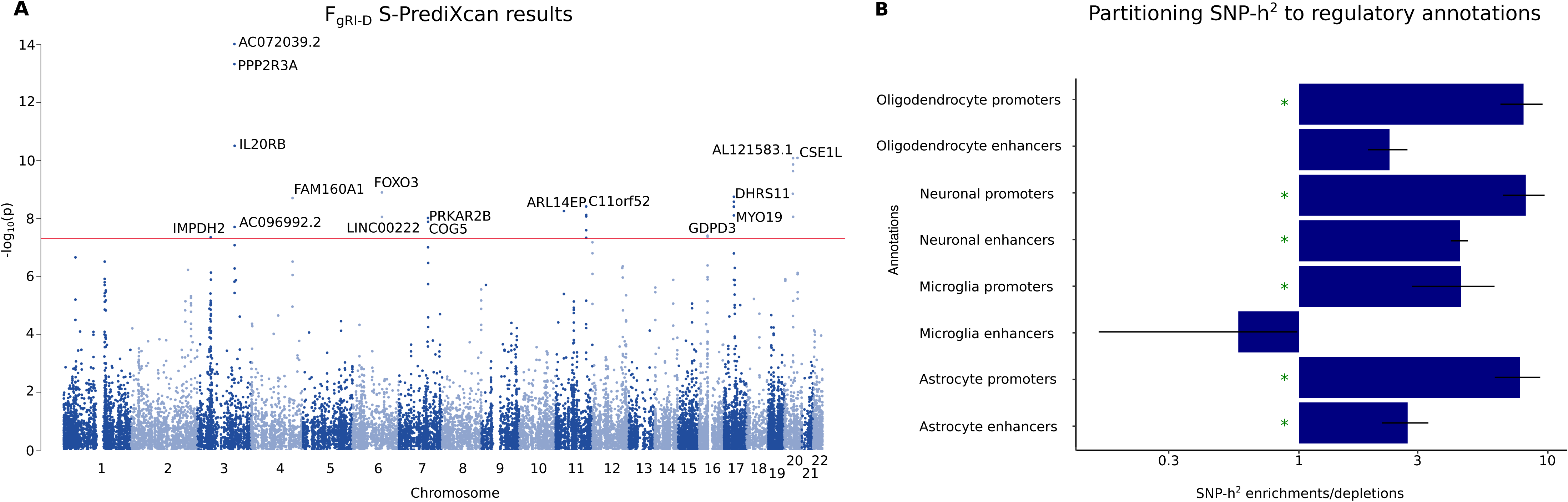
S-PrediXcan and LDSC partitioned heritability results for 8 regulatory brain-cell type annotations. **(A)** Manhattan plot showing TWAS results on 13 brain tissue and whole- blood tissues. Each dot corresponds to a gene-tissue pair. The most significant gene-tissue association pair is shown for each gene. The red line corresponds to the genome-wide significance threshold (P<5×10^−8^). **(B)** Barplots showing LDSC SNP-*h*^2^ enrichment/depletion estimates for each of the 8 regulatory annotations. Green asterisk indicate significance after FDR correction for 8 tests (P_FDR_<0.05). Error bars represent standard errors.

To investigate the neurobiology of genetic variation shared between rhythm impairment and dyslexia at cell-type resolution, we performed LDSC partitioned heritability analysis^35^ using cell-type specific regulatory region annotations of neurons, microglia, astrocytes and oligodendrocytes^36^. We found robust significant SNP-heritability enrichments in the promoters of neurons (Enrichment(SE)=8.14(1.55), P_FDR_=3.38×10^−5^), oligodendrocytes (Enrichment(SE)=7.98(1.53), P_FDR_=3.38×10^−5^), astrocytes (Enrichment(SE)=7.72(1.59), P_FDR_=1.1×10^−4^) and microglia (Enrichment(SE)=4.47(1.63), P_FDR_=0.04), as well as enhancers of neurons (Enrichment(SE)=4.43(0.35), P_FDR_=7.96×10^−18^) and astrocytes (Enrichment(SE)=2.73(0.58), P_FDR_=4.35×10^−3^) (Fig. 3B, Table S12). Consistent with the original rhythm and dyslexia GWAS reports^8,9^, F_gRI-D_ relates to brain structure in part by common effects at regulatory regions within multiple cell-types, including neuronal and various non-neuronal cells such as oligodendrocytes. This may suggest that the F_gRI-D_ might impact myelination and white-matter connectivity patterns that could potentially instantiate neural overlap between rhythm and reading-related aspects of language^1,5,37^.

We then moved on to investigate relationships of F_gRI-D_ with psychiatric, neurological, and behavioural traits, examining patterns of genetic correlations with common and independent factors in more detail. First, we curated 88 sets of GWAS summary statistics including traits that were significantly genetically correlated either with rhythm or dyslexia in the original GWAS reports^8,9^, and three additional education-related traits^19^ (Table S13). To reduce the statistical burden of multiple testing correction in our consequent analyses, we subset this initial set of 88 traits based on their levels of genetic correlation among themselves. To do so, we estimated pairwise genetic correlations, and identified the most highly correlated traits (|*r_g_*|>0.80; Fig. S5). We then performed hierarchical clustering, obtaining one representative trait from each cluster of highly correlated traits (Fig. S6). This approach yielded 49 traits that were relatively genetically independent (see Methods for details), for which we estimated the genetic correlations with F_gRI-D_, and with the summary statistics obtained by the IPM (Fig. S7, Table S14). Genetic correlations between F_gRI-D_ and the assessed traits ranged from −0.56 to 0.46, and mostly lay between the genetic correlation estimates for independent factors (Fig. S7), supporting that F_gRI-D_ indeed captures the common genetic factor of rhythm impairment and dyslexia. We found significant negative correlations between F_gRI-D_ and non-word repetition (*r_g_*(SE)=-0.513(0.099), P_FDR_=7.03×10^−7^), and phoneme awareness (*r_g_*(SE)=-0.562(0.058), P_FDR_=3.78×10^−21^), validating the F_gRI-D_ construct’s link to reading- and language-related traits. Positive genetic correlations were observed for ADHD (*r_g_*(SE)=0.237(0.029), P_FDR_=3.69×10^−15^), autism spectrum disorder (*r_g_*(SE)=0.075(0.035), P_FDR_=4.529×10^−2^), and insomnia (*r_g_*(SE)=0.200(0.027), P_FDR_=6.04×10^−13^), suggesting shared genetic liability with neuropsychiatric traits that have been phenotypically linked to rhythm^38^. In total, F_gRI-D_ showed significant (P_FDR_<0.05) genetic correlations with 37 of the 49 selected psychiatric/neurological/behavioural traits with varying magnitudes and directions, including ADHD, Parkinson’s Disease, health satisfaction and loneliness (|*r_g_|* median=0.146, range=0.06-0.56). Consistent with the ARRH hypothesis, the directionality of genetic correlations suggest that decreased rhythm impairment/dyslexia risk may be associated with resilience to certain neuropsychiatric disorders. These genetic correlations also reflect a shared genomic architecture underlying rhythm, dyslexia and social traits, showing that social function and co-evolution hypotheses of rhythm and communication skills^39,40,41^ are plausible from a genetic perspective. Future work will be needed to disentangle possibly shared genomic substrates of the evolution of social interaction, language and music.

Even though reading is a recent human innovation, it recruits language-related brain circuits^42,43^, which have undergone biological evolution on the lineage leading to humans. Similarly, dyslexia manifests overtly as a reading/spelling disorder, yet in many cases this reflects underlying deficits in aspects of oral language (e.g. phonological awareness)^10,11,12,13^. Given this link between spoken language and reading, and in light of theoretical frameworks positing co-evolution of rhythm- and language-related skills in humans^5,39,40,41,44^, we leveraged genomic methods to investigate the evolution of the overlap between rhythm and the reading-related aspect of language over a range of timescales (Fig. 4A). We first performed LDSC partitioned heritability analysis using five evolutionary annotations tagging foetal brain human gained enhancers^45^, Neandertal introgressed alleles^46^, archaic deserts^47^, and primate conserved and accelerated regions^48^ (Fig. 4A). This revealed significant SNP-heritability depletions in Neandertal introgressed alleles, and significant enrichments in primate conserved regions for all traits (Fig 4B, Table S15), in line with findings for many other complex human traits^49^. We then used the SBayesS function of the GCTB package^50^ to probe the effect size-minor allele frequency relationship (*Ŝ*) – an essential component of the complex trait genetic architecture influenced by natural selection^50^. Similar to most cognitive and behavioural traits^50^, we found moderate levels of negative selection acting on F_gRI-D_ (*Ŝ*(SD)*=*-0.51(0.05)), and the independent factors of dyslexia *Ŝ*(SD)*=*-0.47(0.06)) and rhythm impairment (*Ŝ*(SD)*=*-0.49(0.06)) (Fig. 4D, Table S16). To pin down gene-sets associated with various evolutionary events and timescales that are not testable via partitioned heritability analysis, we performed MAGMA gene-set analysis^51^. Specifically, we tested whether genetic variation associated with F_gRI-D_ was enriched in genes that overlap with four evolutionary annotations (Tables S17-20): i) Ancient Selective Sweep sites^52^, ii) Human Accelerated Regions^53,54,55,56^, iii) Differentially Methylated Regions (DMRs) between Anatomically Modern Humans (AMHs) and archaic humans^57^, and iv) DMRs between AMHs and chimpanzees^57^. These gene-set based analyses did not yield any significant enrichment signals (Table S21), indicating an absence of evidence for associations between F_gRI-D_ and these four annotations.

**Fig. 4:**
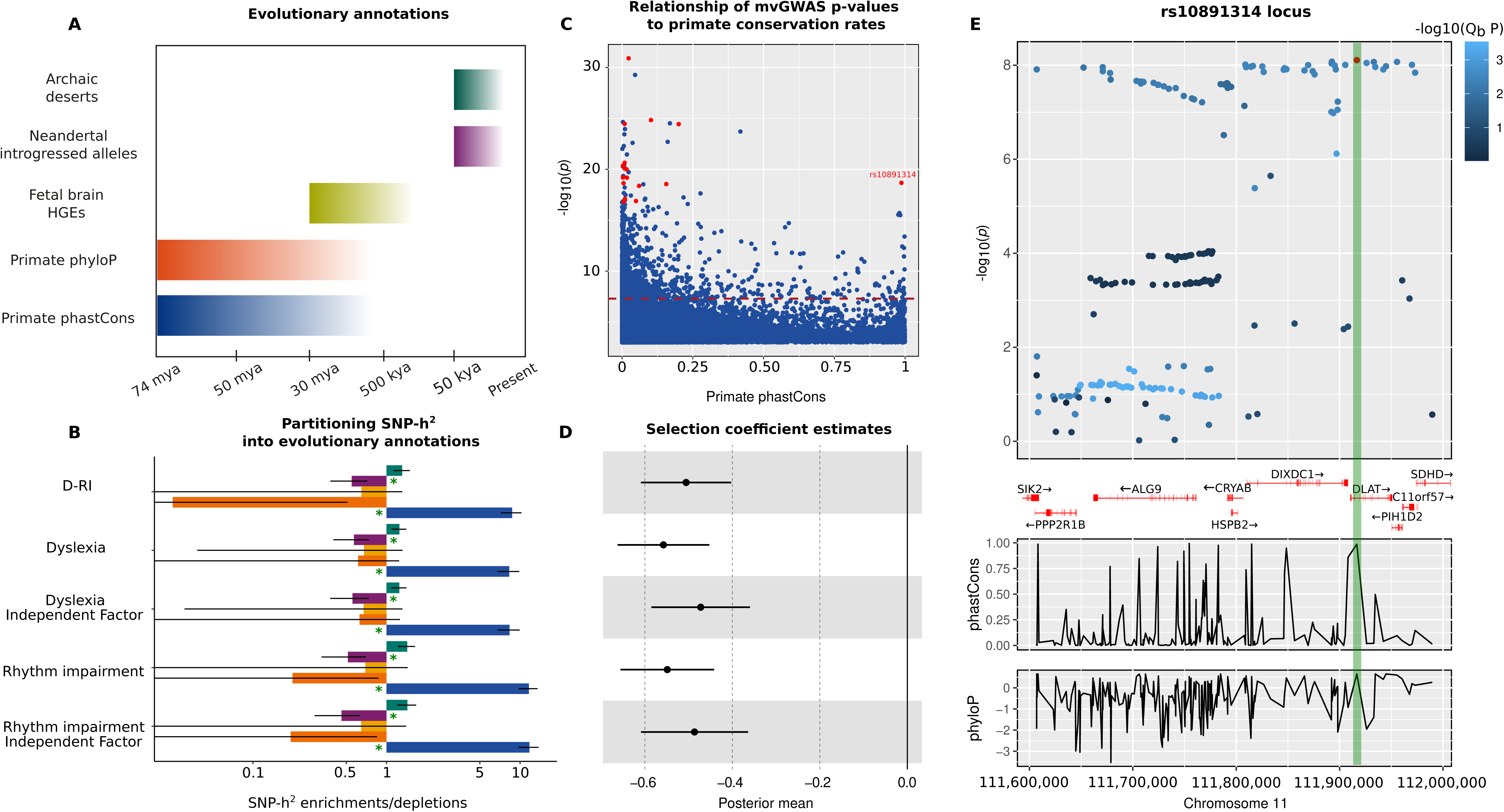
Evolutionary analyses of dyslexia, rhythm impairment, F_gRI-D_ and independent factors. **(A)** Timescales covered by evolutionary annotations that we used. **(B)** LDSC partitioned heritability estimates for each annotation-trait pair. Colour coding of the bars correspond to annotations in panel A. Green asterisk indicate significance after FDR correction for 25 tests (P_FDR_<0.05). Error bars represent standard errors. **(C)** A scatter plot showing the association between F_gRI-D_ mvGWAS -log_10_(P) values and primate phastCons scores. Lead SNPs in 17 genome-wide significant loci are highlighted as red data points (1 missing genome-wide significant locus lead SNP does not have a phastCons score). The dashed red line indicates genome-wide significance threshold (P<5×10^−8^). **(D)** GCTB SBayesS selection coefficient estimates as posterior means. Error bars represent standard errors. **(E)** Results of a manual look- up of the SNP rs10891314, showing its co-localization with *DLAT*. Colour coding reflects Q_b_ scores. PhastCons and phyloP panels below show patterns of primate conservation and accelerated evolution along the haplotype.

To follow up the significant partitioned SNP-heritability enrichments in primate conserved regions, we investigated the association between F_gRI-D_ mvGWAS p-values and per-SNP primate phastCons scores^48^ for 38,164 clumped SNPs (P*<*0.05, *r*^2^<0.06) from F_gRI-D_ summary statistics (Fig. 4C), and found that one of the F_gRI-D_ genome-wide significant hits, rs10891314, had an exceptionally high phastCons score, likely because it is a missense variant (Fig. 4C). We zeroed- in on this genome-wide significant hit as an example locus and dissected patterns of Q_b_, and conservation/accelerated evolution in primates (Fig. 4E), confirming the sharp increase in conservation rate for the SNP rs10891314. The Human Genome Dating Atlas^58^ estimates this polymorphism to be 11,199 generations old (95% confidence interval), corresponding to ∼280,000 years ago assuming 25 years per generation, around the time period when the oldest known *Homo sapiens* fossils have been dated^59^. Rs10891314 is located in the *DLAT* gene, which is associated with a rare neurodevelopmental disorder Pyruvate Dehydrogenase E2 deficiency characterised by neurological dysfunction, dystonia and learning disability mainly appearing during childhood^60^. *DLAT* is highly conserved and loss-of-function intolerant (pLI=6.68)^61^, which makes this particular missense variant an interesting candidate for increasing susceptibility to rhythm impairment and dyslexia.

After assessing evolutionary signatures on F_gRI-D_ at the genome-wide and SNP levels, we extended our investigations of rhythm-language co-evolution by integrating with independent data from neuroimaging genetics. Thus, we estimated local genetic correlations between F_gRI-D_ and fractional anisotropy (FA) measures of five left hemispheric white-matter tracts (Table S22), involved in the dorsal stream of spoken language, and theorized as key components of rhythm- language convergent evolution^5,62^. Using LAVA^63^, we identified a significant genetic correlation between F_gRI-D_ and the left hemispheric superior longitudinal fasciculus (SLF) I (*r_g_*=1, P_FDR_=0.02) (Table S23) on a ∼2mb region on chromosome 20 (30,569,660-32,484,506) which encompasses several genes including *EFCAB8*, *BAK1P1* and *SUN5* (Fig. S8). SLF-I is the dorsal division of SLF connecting the superior parietal and superior frontal lobes^64^, with functional links to musical rhythm^65^. This finding is consistent with the hypothesized role of the dorsal stream in supporting co-evolution of phonological processing and beat synchronisation^4^.

In summary, we showed robust genetic correlations between rhythm and a number of reading- and language-related traits, supporting ARRH. The bivariate Genomic SEM approach that we developed allowed us to identify genetic overlaps between rhythm impairment and dyslexia, and to present a map of homogeneous and heterogeneous genetic effects, shedding light on patterns of pleiotropy between the two. Our post-mvGWAS analyses enhance our understanding of the aetiology of rhythm and language (on which reading depends) by revealing intricate links across rhythm impairment, dyslexia, and various aspects of evolutionary past and neurobiological function (including gene expression in brain tissue, brain cell type-specific gene regulation, and a local genetic correlation with a tract linked to processing and production of speech and music) ^5^. The evolutionary analyses aimed to provide empirical genetic data as groundwork towards understanding potential evolutionary forces acting jointly on human rhythm- and language-related skills^44,66^, revealing a candidate gene, *DLAT*, for future experimental investigations.

Despite a number of practical constraints, such as the fact that the source GWASs were performed in European-only cohorts, and potential confounds due to residual population stratification and socioeconomic factors, our study represents a first step towards characterising the shared genetic architecture between rhythm- and language-related traits. We reveal complex links across common DNA variants, genes, genomic loci, white-matter structures and human behaviour, making a first set of links across the immensely long causal chain spanning these layers. Developing and applying more sophisticated methods to dissociate environmental confounds from genetics will allow future studies to obtain a better understanding of the genetics and evolution of human language and musicality.

## Methods

### GWAS summary statistics

Beat synchronisation and dyslexia GWAS summary statistics^8,9^ were obtained from 23andMe Inc., a customer genetics company. Both GWASs were performed on European ancestry individuals through online participation and participants provided informed consent. The 23AndMe sample prevalence of dyslexia is 4.6% (N_total_=1,138,870, mean age=51), and sample prevalence of beat synchronisation is 92% (N_total_=606,825, mean age=52). Summary statistics files were reformatted and harmonised to include required columns (e.g. SNP ID, beta, beta S.E., p-value) for each mvGWAS tool following the guidelines in original publications of each tool. To obtain rhythm impairment summary statistics, effect sizes in the binary beat synchronisation GWAS summary statistics were multiplied by -1, so that the effect directions were reversed. Yielding set of GWAS summary statistics comprised of SNP effects contributing to rhythm impairment, and was used for the subsequent mvGWAS analysis with dyslexia. We applied GC correction to both sets of summary statistics for all non-LDSC-based analyses. For LDSC-based analyses (including Genomic SEM), uncorrected summary statistics were used as input, as GC correction biases the LDSC SNP-heritability estimates downwards. The resulting set of summary statistics from Genomic SEM was GC corrected.

### SNP-heritability and genetic correlation estimations

We used LDSC^16^ (v1.0.1) to estimate the SNP-heritabilities and genetic correlations. For rhythm impairment and dyslexia, we estimated the total SNP-heritability on a liability scale using population and sample prevalence information from the original studies (sample prevalence of 0.045 for dyslexia and 0.085 for rhythm impairment, and a population prevalence of 0.050 for dyslexia and 0.048 for rhythm impairment). Genetic correlations were estimated using bivariate LDSC between rhythm, dyslexia, GenLang quantitative reading-/language-related traits^18^, Danish School Grades GWAS^19^, and all external summary statistics except for the planum temporale asymmetry and the language resting-state functional connectivity, which were assessed as described below.

To estimate genetic correlations between rhythm and planum temporale asymmetry^21^, and between rhythm and language resting-state functional connectivity^20^, we used an approach proposed by Naqvi et al.^67^ applicable to unsigned multivariate statistics, as the mvGWAS effect sizes or beta values, which are required to run genetic correlation analysis using LDSC, were not available for these traits. We evaluated the amount of shared signal between each pair of GWASs by estimating the Spearman correlation of the average SNP p-values within approximately independent LD blocks^68^. We first filtered the genome-wide SNPs using the HapMap3 reference panel without the MHC region (https://github.com/bulik/ldsc). We then split the genome-wide SNPs into 1,703 approximately independent blocks^68^. For each approximately independent LD block, we computed the average SNP −log_10_(p-value). We then estimated a rank-based Spearman correlation using the averaged association value (n=1,703) for each LD block. A standard error of the Spearman correlation was estimated using statistical resampling with 10,000 bootstrap cycles with replacement from the 1,703 LD blocks.

### Multivariate genome-wide association studies

To investigate the shared genetic variance of rhythm impairment and dyslexia, we performed multivariate GWASs using three tools: Genomic SEM^17^, N-weighted GWAMA^24^, CPASSOC^25^. These tools use GWAS summary-level data and account for genetic correlation and sample overlap using the cross-trait LD score regression intercept.

#### Genomic SEM (Common and Independent Pathway Models)

First, we reformatted our summary statistics for LDSC (munged) and Genomic SEM, following standard guidelines (https://github.com/GenomicSEM/GenomicSEM/wiki). We then used the multivariable extension of LDSC to estimate the 2*×*2 empirical genetic covariance matrix between rhythm impairment and dyslexia and their associated sampling covariance matrix. We specified a measurement model (Fig. S1), where a shared genetic factor (F*_g_*) was estimated to capture the observed genetic covariance between rhythm impairment and dyslexia. Given that the number of observed parameters for any 2*×*2 covariance matrix equals 3, we constrained all paths between F*_g_* to 1. The final Common Pathway Structural model (CPM) was fit to a genetic covariance matrix which incorporates the SNP tested (Fig. S1), SNPs were regressed from F*_g_*, and residuals were freely estimated. The 1000 Genomes Phase 3 reference panel^69^ was used as the reference panel to calculate SNP variance across traits. Effective population size per-GWAS was calculated as 4×N_cases_×(1-N_cases_/N_total_). Both the reference panel and effective population sizes were then fed into the sumstats function and summary statistics were prepared for the meta-analysis. We applied genomic correction to the CPM results based on the genomic inflation index estimated by LDSC (λ_GC_=1.62; Fig. S2). The final Independent Pathways model (IPM), was fit to the same matrices incorporating the SNP effects, but with the SNP effect being directly regressed from the traits. The final bivariate heterogeneity score, Q_b,_ was obtained by subtracting by a χ^2^ difference test, where the χ^2^ of the IPM is subtracted from the χ^2^ of the CPM (Q_b_ = χ^2^ – χ^2^)^22^. High Q_b_ value index that the association between the SNP and rhythm impairment or dyslexia is not well accounted for by the factor F_g_. We then used the intersect function of bedtools (v. 2.29.2)^70^ to identify the overlaps between genome-wide significant Q_b_ (Table S3), and CPM loci (Table S2), as well as +-1Mb surroundings of each CPM locus.

#### CPASSOC

Following the CPASSOC manual^25^, we used the median sample size for each summary statistics file as 23andMe SNPs can have varying sample sizes. We removed SNPs with a Z-score larger than 1.96 or less than -1.96, and extracted a 2×2 genetic correlation matrix for dyslexia and rhythm impairment. Then we generated a *M*×*K* matrix of summary statistics where each row represented a SNP, and 2 columns represented dyslexia and rhythm impairment Z-scores. We finally performed the S_hom_ test, and obtained a vector of p-values for *M* SNPs using pchisq function in R (4.0.3).

#### GWAMA (N-weighted)

To account for sample overlap, we first generated a matrix of cross-trait intercepts using the intercepts of LDSC genetic correlations between dyslexia and rhythm impairment summary statistics. We then performed N-weighted GWAMA by feeding the Cross Trait Intercept matrix and a vector of SNP-heritabilities of each trait using the multivariate_GWAMA function.

### Transcriptome-wide association study

We conducted a transcriptome-wide association study (TWAS) using S-PrediXcan framework^28^ and the joint-tissue imputation (JTI) TWAS derived models from GTEx v8 tissues^21^. PrediXcan predicts gene expression from the genotype profile of each individual by using the JTI model weights, which were trained on GTEx^71^, and validated on PsychEncode^72^ and GEUVADIS^73^. These SNP-expression weights represent the correlations between SNPs and gene expression levels. To overcome the requirement for individual-level genotype data, Barbeira et al.^28^, derived a mathematical expression, implemented in S-PrediXcan framework, which effectively yields similar outcomes to PrediXcan using GWAS summary statistics. S-PrediXcan and JTI weights account for LD and collinearity problems due to high expression correlation across tissues^21^. We filtered the 17q21.31 inversion region (∼1.5 Mb long), which has multiple phenotypic associations with brain-related traits^74^ to minimise the impact of this high-LD region on our results. We then corrected TWAS p-values for 192,905 gene-tissue pairs, and used Z-scores and P_FDR_ of the significant (P_FDR_<0.05) pairs to assess gene-F_gRI-D_ associations.

### Gene-set enrichment and pathway analyses

We used PANTHER to run statistical overrepresentation analysis in 3 Gene Ontology (GO) and 3 PANTHER GO-Slim terms (biological process, molecular function, cellular component)^29,30,31,32^ with 315 unique genes that we obtained from TWAS. We used 20,102 genes that we tested in TWAS as the background gene set. Results were FDR corrected for all GO and GO-Slim terms (n=15,028).

### LDSC partitioned heritability with cell type-specific annotations

We used 8 human genome annotations by Nott et al.^36^ tagging promoter and enhancer regions of neurons, oligodendrocytes, microglia and astrocytes using LDSC partitioned heritability analysis^35^ following the guidelines in the LDSC Wiki page (https://github.com/bulik/ldsc/wiki/Partitioned-Heritability). All enrichment analyses were controlled for the baselineLD model v2.2. Enrichment p-value results were FDR corrected for 8 tests.

### Genetic correlations using GWAS summary statistics from neuropsychiatric/behavioural phenotypes

We first compiled 88 traits that were significantly genetically correlated either with rhythm impairment or dyslexia in the original respective GWAS papers^8,9^. We filtered these traits in order to avoid unnecessary multiple testing burden and to focus on genetically independent phenotypes. We first identified 46 traits that are more than +/-80% genetically correlated with at least one other trait. Then we created a distance matrix from the correlation estimates and performed hierarchical clustering using Ward’s method^75^ as the linkage method, which maximises the within-cluster homogeneity to identify trait clusters. We identified 7 clusters using the so-called elbow method, and chose the most informative and representative trait for each cluster based on the highest correlation between traits and the cluster principal component. We added these 7 cluster-representative traits to the remaining 42 traits and used LDSC to estimate genetic correlations with F_gRI-D_ and 2 independent factors. Genetic correlation p-values were FDR corrected for 49 tests.

### Partitioned heritability analysis with custom evolutionary annotations

We used LDSC^16^ (v1.0.1) to estimate partitioned SNP-heritability enrichments/depletions in foetal brain human-gained enhancers, Neandertal introgressed alleles, archaic deserts, conserved loci in the primate phylogeny (Conserved_Primate_phastCons46way annotation from baselineLD), and genomic loci that have a primate phyloP score^48^ less than -2 (presumably suggesting accelerated evolution). All annotations were controlled for baselineLD model v2.2. Foetal brain human-gained enhancers were also controlled for foetal brain active regulatory elements from the Roadmap Epigenomics Consortium database^76^.

### MAGMA gene-set analysis with custom evolutionary gene lists

We compiled four additional evolutionary genomic annotations for MAGMA gene-set analysis^51^ which cover timescales from ∼6 million years ago to ∼250 thousand years ago: Ancient Selective Sweeps^52^, Human Accelerated Regions^53,54,55,56^, Anatomically Modern Human-derived DMRs^57^, and Human vs. chimpanzee DMRs^57^. These annotations either tag regulatory or selective sweep sites. We listed the genes that fall within +/-1 kilobase of each locus tagged by each annotation, and filtered these initial gene lists for protein-coding genes using NCBI’s hg19 genome annotation^77^. The resulting protein-coding gene lists were used for MAGMA gene-set enrichment analysis for rhythm impairment, dyslexia and F_gRI-D_ summary statistics. We first performed gene annotation by integrating SNP locations from the summary statistics, and gene locations from NCBI hg19 genome annotation. We then performed a gene analysis using SNP p-values and 1000 Genomes Phase 3 European panel^69^. We finally applied a gene-set analysis using results from gene annotation and gene analysis, and four gene-sets. Enrichment p-values were FDR corrected for four tests.

### Genome-wide negative selection estimation

We performed SBayesS analysis on the rhythm impairment, dyslexia, F_gRI-D_, and two independent factor GWAS summary statistics using the GCTB software (version 2.02)^50^ to quantify the level of negative selection acting on these traits. SBayesS estimates total SNP-heritability, polygenicity, and the relationship between variants’ minor allele frequencies and effect sizes, and generates a genome-wide negative selection metric (*S*) which ranges from 0 to -1. *S* estimates that are closer to -1 are interpreted as a sign of strong negative selection^50^, whereas estimates closer to 0 can suggest positive selection (see Zeng et al., 2021).

### LAVA local genetic correlations with white-matter connectivity measures

To identify local regions of the genome that might be shared between rhythm, language and evolutionarily relevant brain circuitry, we tested local genetic correlations between F_gRI-D_ and white-matter connectivity measures. We performed GWASs of selected brain imaging traits using data from the UK Biobank^78^. For these GWASs, UK Biobank data first underwent sample and genetic quality control and brain imaging data processing, followed by genome-wide association analysis.

#### Sample quality control

This study used the UK Biobank February 2020 release (research application number: 79683). All participants provided informed consent and the study was approved by the North West Multi-Center Research Ethics Committee (MREC). For individual with both diffusion-weighted MRI and genotyping data, we excluded participants with unusual heterozygosity (principal components corrected heterozygosity>0.19), high missingness (missing rate>0.05), sex mismatches between genetically inferred sex and self-reported sex as reported by Bycroft et al.^78^. We further restricted our analyses to individuals with white British ancestry as defined by Bycroft et al.^78^ in order to avoid any possible confounding effects related to ancestry. This resulted in 31,465 individuals (mean age=55.21 years old, range between 40 to 70 years old, 16,497 females) passing the sample QC.

#### Genetic quality control

The imputed genotypes were obtained from the UK Biobank portal. These data underwent a stringent quality control protocol. We excluded SNPs with minor allele frequencies below 1%, Hardy-Weinberg p-value below 1×10^−7^ or imputation quality INFO scores below 0.8. Multiallelic variants which cannot be handled by many programs used in genetic-related analyses were removed. This resulted in 9,422,496 autosomal SNPs that were analyzed in the GWAS.

#### Neuroimaging phenotypes

Neuroimaging measures of white-matter tracts were derived from the diffusion-weighted scans (3T Siemens Skyra scanner) released by the UK Biobank Imaging Study (refer to http://biobank.ndph.ox.ac.uk/showcase/refer.cgi?id=2367 for the full protocol). Briefly, in vivo, whole-brain diffusion-weighted MRI scans were acquired and fed into Diffusion Tensor Imaging (DTI) modelling to assess brain microstructure and derive a fractional anisotropy (FA) quantitative diffusion map that was subject to a TBSS (tract-based spatial statistics) analysis resulting in a skeletonised image. Details of the image acquisition, quality control and processing are described elsewhere^79^. We extracted the following regions: The left arcuate fasciculus (long, anterior, and posterior segments), the left superior longitudinal fasciculus (I, II, III), and the left uncinate fasciculus for each individual by averaging the FA skeletonised image across a set of five left white-matter tracts defined from a probabilistic atlas^80^. *Genome-wide association scanning.* GWASs were performed separately for each of the neuroimaging phenotypes using imputed genotyping data, with PLINK (v1.9)^81^. We made use of categorical and continuous variables controlling for covariates in the GWASs including age, sex, genotype array type, and assessment centre. To avoid possible confounding effects related to ancestry, we used the first ten genetic principal components capturing population genetic diversity. These covariates are considered in a pre-residualization step: a multiple linear regression of the endophenotype vector on the covariates is performed and all these ones are replaced by their corresponding residual. Additionally, a rank-based inverse normalization is performed to ensure that the distributions of endophenotypes are normally distributed.

### Local genetic correlations

We identified a list of overlapping loci using 2,495 LD blocks covering the whole human genome provided in the Local Analysis of [co]Variant Association (LAVA)^63^ partitioning algorithm GitHub repository (https://github.com/cadeleeuw/lava-partitioning), and 1,609 genome-wide significant (P<5×10^−8^) SNPs in our F_gRI-D_ summary statistics. This resulted in 18 LD blocks. We then used LAVA to estimate local genetic correlations between F_gRI-D_ and the five aforementioned white-matter tracts. LAVA estimates local heritability for each of these 18 LD blocks, and for each considered trait. For the loci which explained a significant proportion (nominally significant SNP-heritability estimate, P<0.05) of the total SNP-heritability of F_gRI-D_ and white-matter tracts, we proceeded to perform bivariate local genetic correlation. This extra step of filtering based on local SNP-heritability estimates is not mandatory but recommended^63^. Finally, we obtained local genetic correlation estimates and associated p-values, which we FDR corrected for 14 tests.

## Data availability

The full GWAS summary statistics from the original 23andMe discovery studies set have been made available through 23andMe to qualified researchers under an agreement with 23andMe that protects the privacy of the 23andMe participants. Datasets will be made available at no cost for academic use. Please visit https://research.23andme.com/collaborate/#dataset-access/ for more information and to apply to access the data. Participants provided informed consent and volunteered to participate in the research online, under a protocol approved by the external AAHRPP-accredited IRB, Ethical & Independent (E&I) Review Services. As of 2022, E&I Review Services is part of Salus IRB (https://www.versiticlinicaltrials.org/salusirb).

## Code availability

All scripts used for analyses are publicly available on the GitHub repository: https://github.com/galagoz/pleiotropyevo

This study used openly available software, specifically PLINK (http://zzz.bwh.harvard.edu/plink/), and S-PrediXcan (https://github.com/hakyimlab/MetaXcan). JTI-TWAS prediction models trained on GTEx v8 are available at the PredictDB website (http://predictdb.org) and (https://github.com/gamazonlab/MR-JTI/tree/master). The human frontal lobe probabilistic atlas used is available at (http://www.bcblab.com/BCB/Atlas_of_Human_Brain_Connections.html).

## Acknowledgements

This project was supported in part by funding from the National Institute on Deafness and Other Communication Disorders, the Office of Behavioral and Social Sciences Research, and the Office of the Director of the National Institutes of Health under Award Numbers R01DC016977, K18DC017383, and DP2HD098859. G.A., E.E., G.B., and S.E.F. are supported by the Max Planck Society. G.B. is also supported by the German Federal Ministry of Education and Research (BMBF). The funders had no role in study design, data collection and analysis, the decision to publish, or the preparation of the manuscript. S.E.F. is a member of the Center for Academic Research and Training in Anthropogeny (CARTA). This research was conducted using the UK Biobank resource under application no. 79683. We would like to thank the research participants and employees of 23andMe for making this work possible.

The following members of the 23andMe Research Team contributed to this study: Stella Aslibekyan, Adam Auton, Elizabeth Babalola, Robert K. Bell, Jessica Bielenberg, Jonathan Bowes, Katarzyna Bryc, Ninad S. Chaudhary, Daniella Coker, Sayantan Das, Emily DelloRusso, Sarah L. Elson, Nicholas Eriksson, Teresa Filshtein, Pierre Fontanillas, Will Freyman, Zach Fuller, Chris German, Julie M. Granka, Karl Heilbron, Alejandro Hernandez, Barry Hicks, David A. Hinds, Ethan M. Jewett, Yunxuan Jiang, Katelyn Kukar, Alan Kwong, Yanyu Liang, Keng-Han Lin, Bianca A. Llamas, Matthew H. McIntyre, Steven J. Micheletti, Meghan E. Moreno, Priyanka Nandakumar, Dominique T. Nguyen, Jared O’Connell, Aaron A. Petrakovitz, G. David Poznik, Alexandra Reynoso, Shubham Saini, Morgan Schumacher, Leah Selcer, Anjali J. Shastri, Janie F. Shelton, Jingchunzi Shi, Suyash Shringarpure, Qiaojuan Jane Su, Susana A. Tat, Vinh Tran, Joyce Y. Tung, Xin Wang, Wei Wang, Catherine H. Weldon, Peter Wilton, Corinna D. Wong.

## Author contributions

G.A., E.E., N.J.C., R.L.G., and S.E.F. designed research; G.A., E.E., Y.M., G.B. and E.E. performed research; G.A., E.E., and Y.M. analyzed data; G.A. wrote the initial draft of the manuscript; E.E., Y.M., G.B., P.F., M.G.N., M.L., R.L.G, and S.E.F. provided critical feedback and commented on the manuscript. P.F. is employed by and hold stock or stock options in 23andMe, Inc.

