## Supplementary Information for "The shared genetic architecture and evolution of human language and musical rhythm"

This PDF file includes:
Supplementary Figures 1 to 8

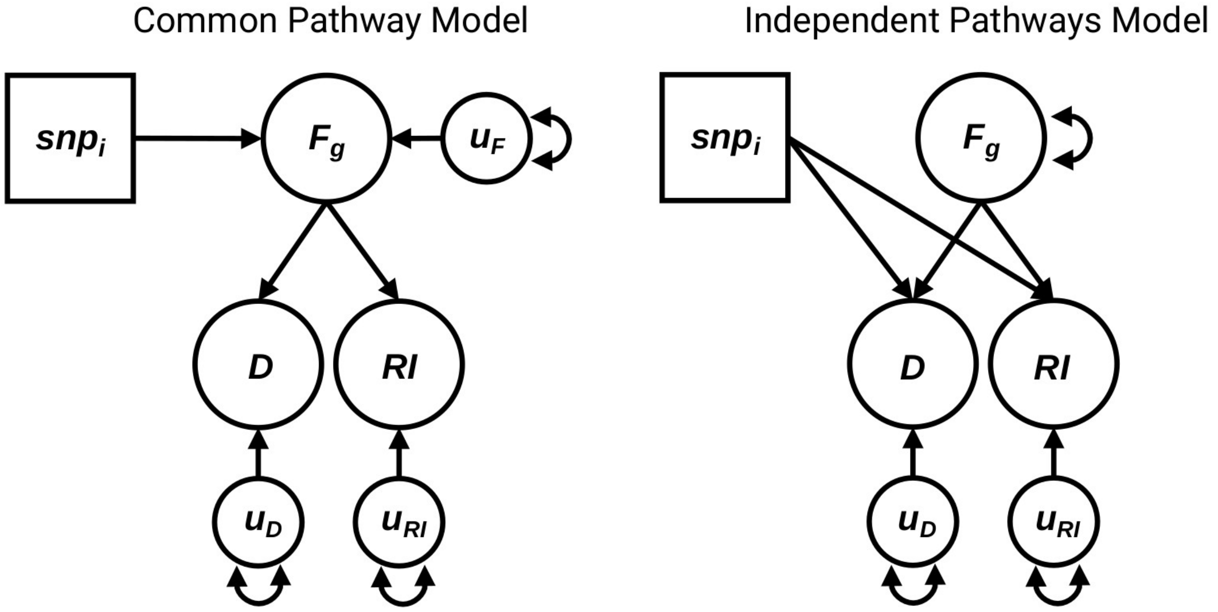


**Supplementary Figure 1:** Genomic SEM model diagrams for common pathway and independent pathways models. The models were used in the mvGWAS for the common factor and independent factor association analyses. D: dyslexia, RI: rhythm impairment, $F_g$: Common factor, $u_D$: residual variance of dyslexia, $u_{RI}$: residual variance of rhythm impairment, $u_F$: residual variance of the common factor, $snp_i$: $i^{th}$ SNP regression.

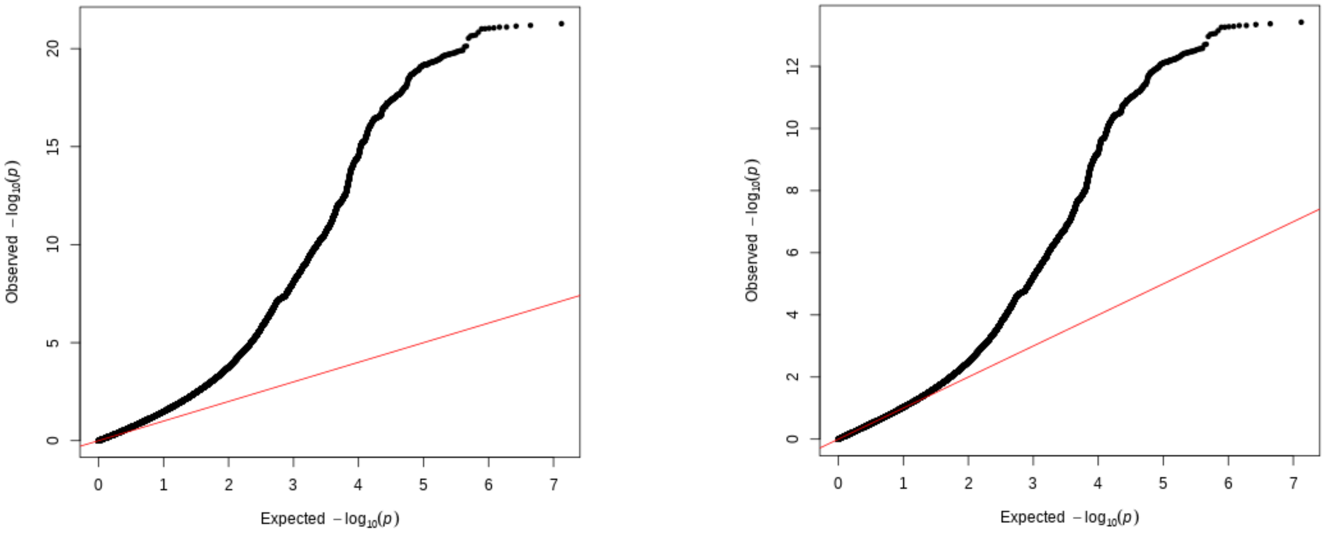


**Supplementary Figure 2:** QQ plots of $F_{gRI-D}$ mvGWAS summary statistics *prior to* (left) and *after* (right) GC correction.

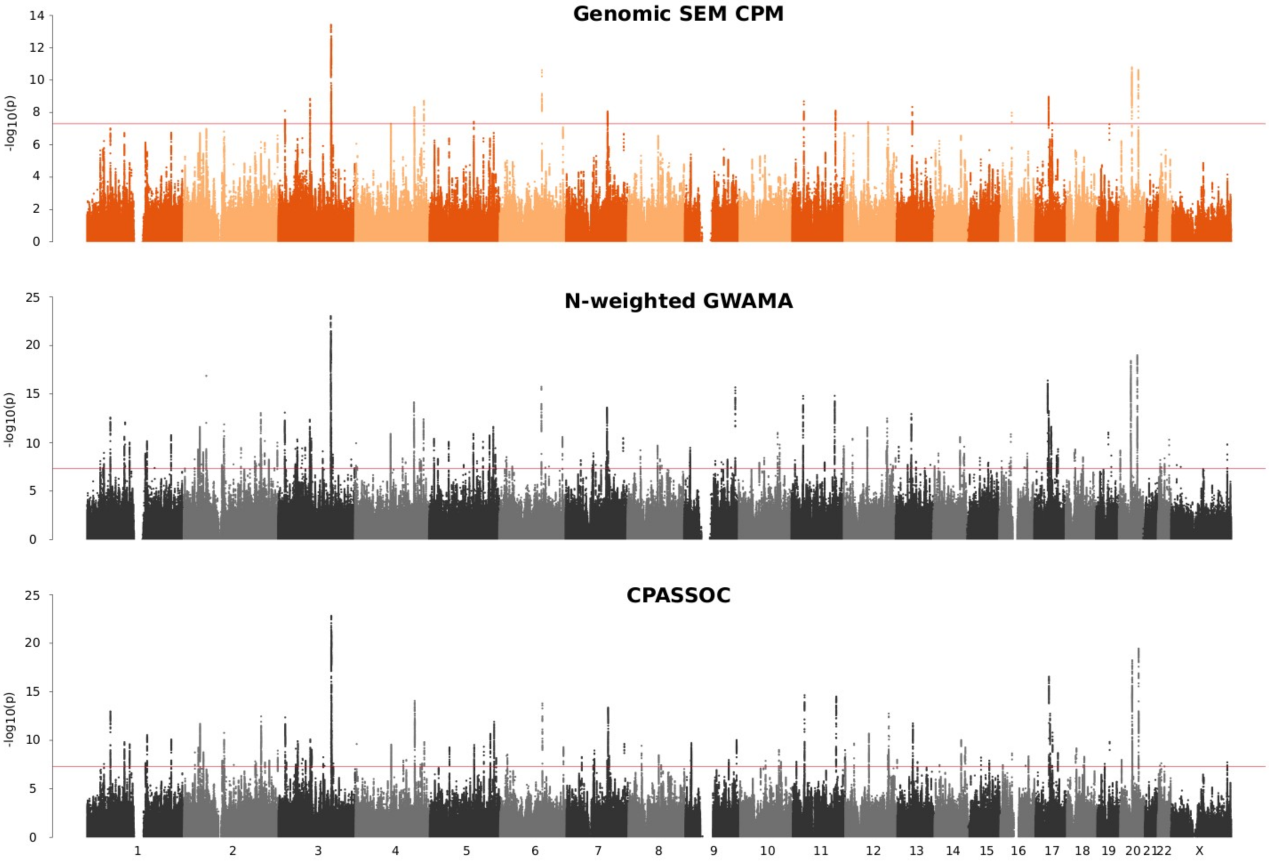


**Supplementary Figure 3:** Manhattan plots for Genomic SEM CPM ($F_{gRI-D}$), N-weighted GWAMA, and CPASSOC. The red lines correspond to the genome-wide significance threshold ($P<5\times10^{-8}$).

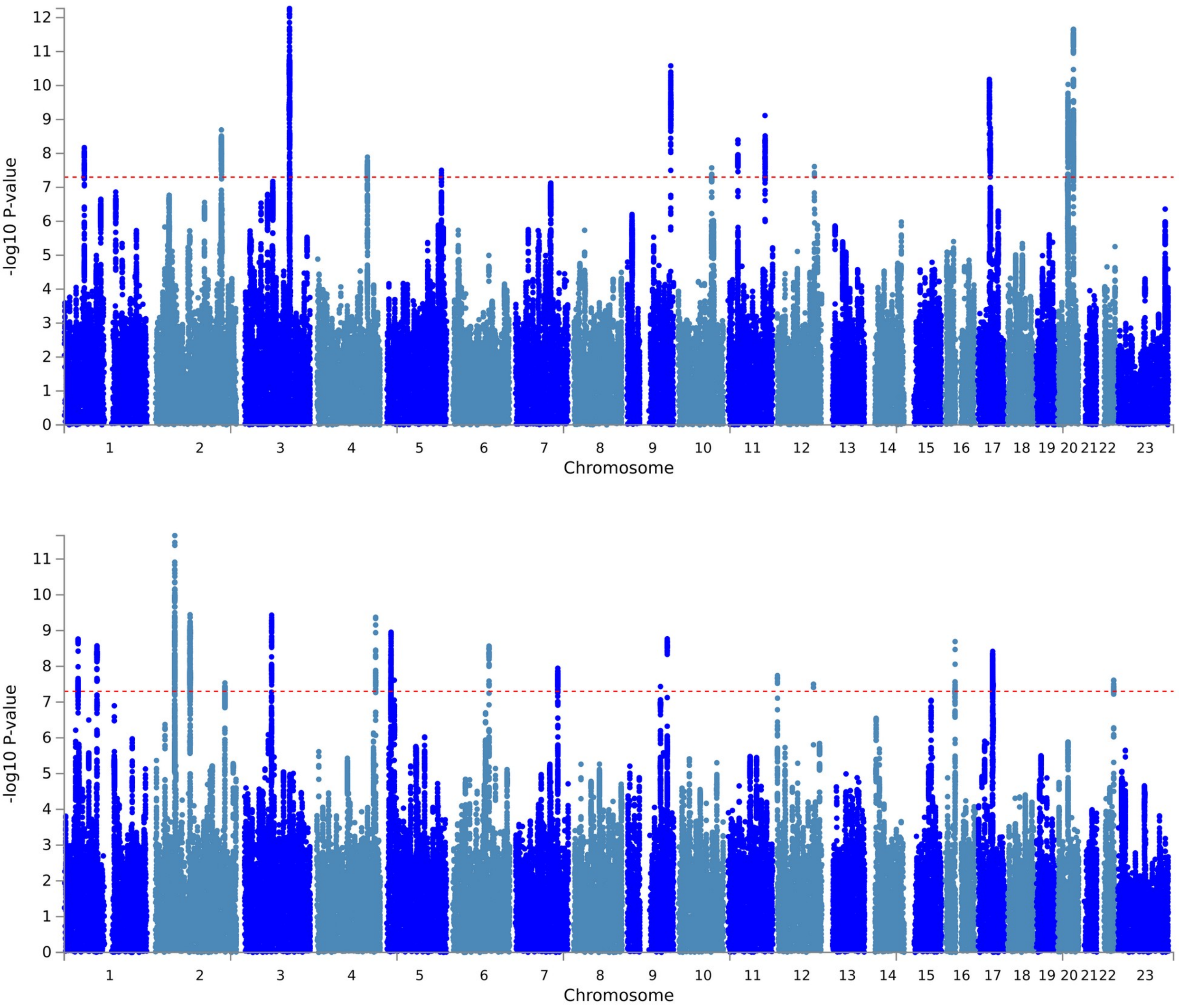


**Supplementary Figure 4:** Manhattan plots for the independent factors of dyslexia (top) and rhythm impairment (bottom). The red lines correspond to genome-wide significance threshold ($P<5\times10^{-8}$).

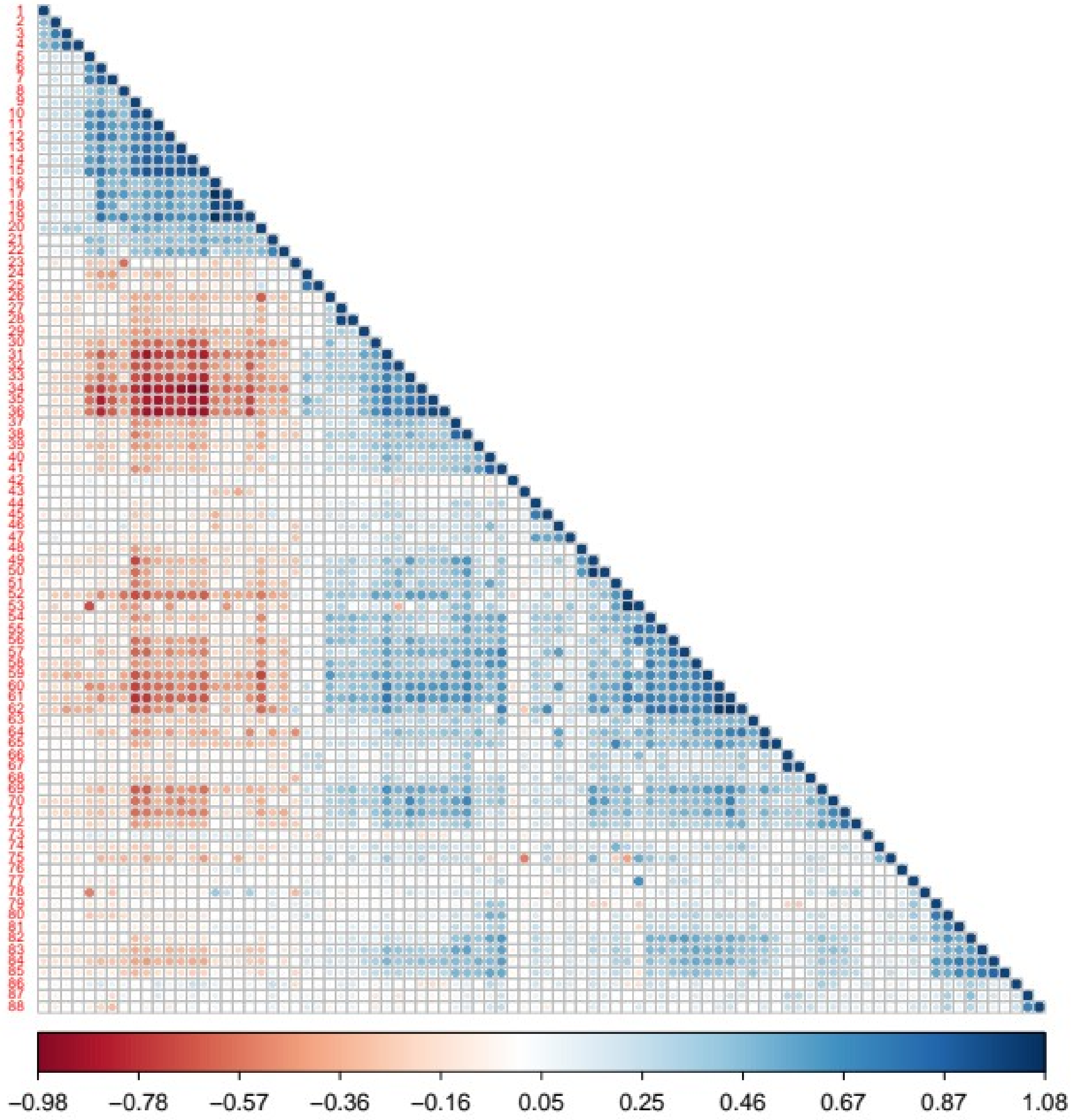


**Supplementary Figure 5:** Genetic correlations ($r_g$) among 88 traits that are significantly correlated either with dyslexia or rhythm. Genetic correlations were estimated using LDSC. Supplementary Table 13 provides the list of traits included.

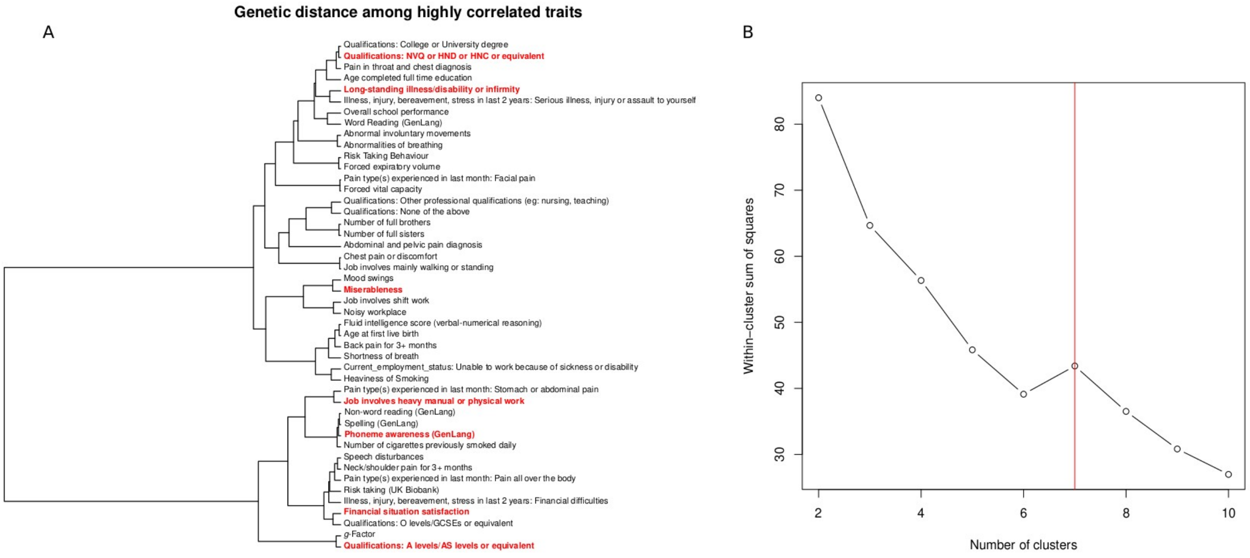


**Supplementary Figure 6:** (A) Dendrogram showing the hierarchical clustering of highly genetically correlated ($|r_g|$>0.80) traits. (B) Knee-point algorithm identified 7 representative clusters with. For each cluster, one representative trait (shown in bold red) was used in the genetic correlation analysis with the $F_{gRI\text{-}D}$.

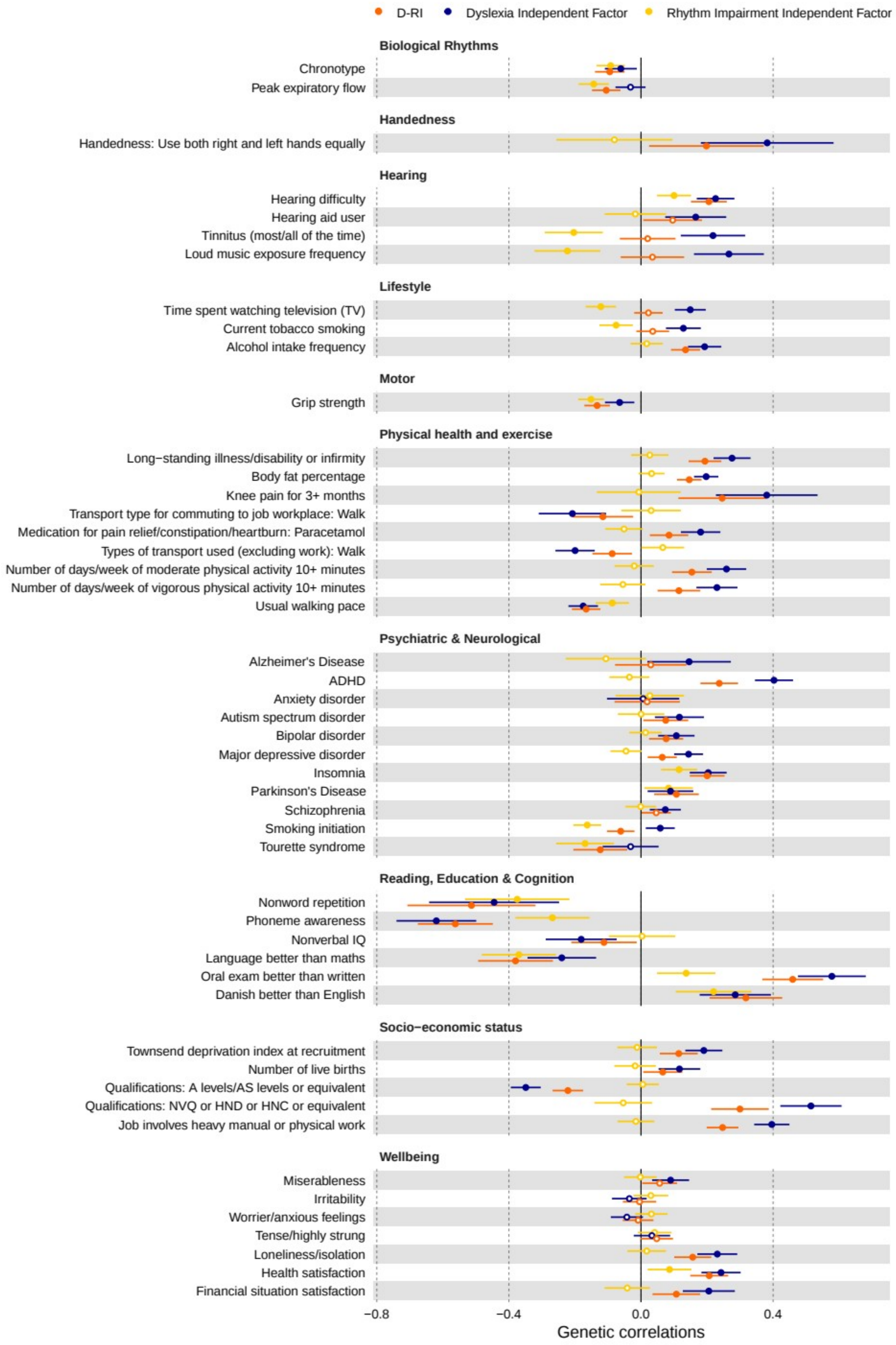


**Supplementary Figure 7:** Genetic correlations ($r_g$) between the selected 49 traits and $F_{gRI\text{-}D}$ (D-RI) (orange) and dyslexia (blue) and rhythm (yellow) independent factors. Genetic correlations were estimated using LDSC. Full circles indicate significant correlations (P<0.05). Error bars represent standard errors.

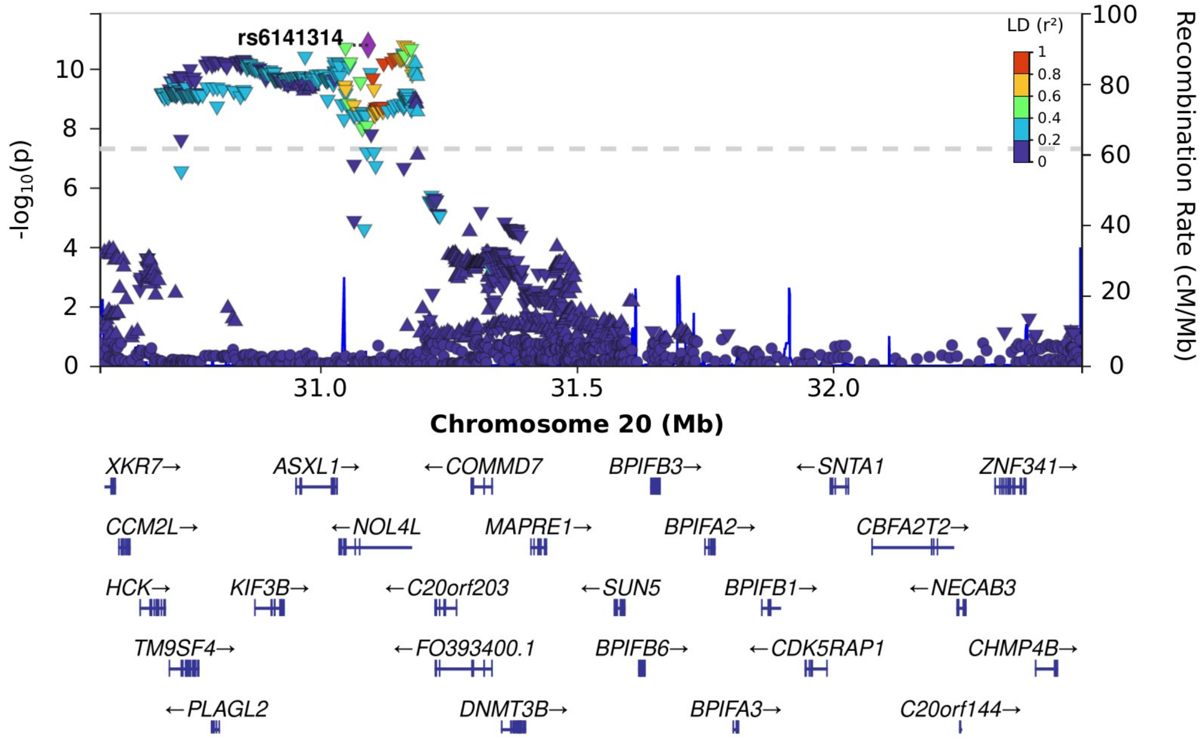


**Supplementary Figure 8:** LocusZoom plot of chr20: 30,569,660-32,484,506 locus, the region which is identified by local genetic correlation analysis of $F_{gRI-D}$ and Superior Longitudinal Fasciculus I. P-values represent $F_{gRI-D}$ mvGWAS significance. Direction of the triangles represent effect directions. LD ($r^2$) levels with rs6141314 are represented in colours. Grey dash-line indicate genome-wide significance level ($P<5x10^{-8}$).
